# Bone mechanoregulation allows subject-specific load estimation based on time-lapsed micro-CT and HR-pQCT *in vivo*

**DOI:** 10.1101/2021.02.22.431960

**Authors:** Matthias Walle, Francisco C. Marques, Nicholas Ohs, Michael Blauth, Ralph Müller, Caitlyn J. Collins

## Abstract

Patients at high risk of fracture due to metabolic diseases frequently undergo long-term antiresorptive therapy. However, in some patients treatment is unsuccessful in preventing fractures or causes severe adverse health outcomes. Understanding load driven bone remodelling, i.e. mechanoregulation, is critical to understand which patients are at risk for progressive bone degeneration and may enable better patient selection or adaptive therapeutic intervention strategies. Bone microarchitecture assessment using high-resolution peripheral quantitative computed tomography (HR-pQCT) combined with computed mechanical loads has successfully been used to investigate bone mechanoregulation at the trabecular level. To obtain the required mechanical loads that induce local variances in mechanical strain and cause bone remodelling, estimation of physiological loading is essential. Yet, current models homogenise strain patterns throughout the bone to estimate load distribution *in vivo*, which may be a flawed assumption for investigating alterations in bone mechanoregulation. By further utilising available spatiotemporal information of time-lapsed bone imaging studies, we developed a mechanoregulation-based load estimation algorithm (MR). MR calculates organ scale loads by scaling and superimposing a set of predefined independent unit loads to optimise measured bone formation in high, quiescence in medium and resorption in low strain regions. We benchmarked our algorithm against a previously published load history algorithm (LH) using synthetic data, micro-CT images of murine vertebrae under defined experimental *in vivo* loadings and HR-pQCT images from seven patients. Our algorithm consistently outperformed LH in all three datasets. *In silico* generated time evolutions of distal radius geometries (n = 5) indicated significantly higher sensitivity, specificity, and accuracy for MR than LH (p < 0.01). This increased performance led to substantially better discrimination between physiological and extra-physiological loading in mice (n = 8). Moreover, a significantly (p < 0.01) higher association between remodelling events and computed local mechanical signals were found using MR (CCR = 0.42) than LH (CCR = 0.38) to estimate human distal radius loading. Future applications of MR may enable clinicians to link subtle changes in bone strength to changes in day-to-day loading, identifying weak spots in the bone micro-structure for local intervention and personalised treatment approaches.

## Introduction

Considerable patient variability poses a severe problem in the clinical assessment and treatment of metabolic bone diseases such as osteoporosis. Diagnosis and bone strength assessment rely heavily on radiographic measures of bone mineral density (BMD). However, sources of error in BMD measurements, i.e. intra- and interpatient variability, makes it challenging to attribute measured BMD changes to the actual biological change (Nguyen et al., 1997). Accordingly, the sensitivity and specificity of predicting individual patients’ risk for fracture are low (Trémollieres et al., 2010; Cervinka et al., 2017). As a consequence, patients may receive treatment, although only a minority would have suffered from a bone fracture. This is relevant because anti-resorptive therapies can result in severe adverse events, including osteonecrosis, hypocalcaemia and thromboembolism (Chen and Sambrook, 2012). Moreover, current diagnostic approaches fail to identify the specific weak spots in the bone. Therefore, they do not estimate where and how fractures will occur and how a local intervention could prevent them (Schultz and Wolf, 2019).

High-resolution peripheral quantitative computed tomography (HR-pQCT), an emerging diagnostic modality of the peripheral skeleton, allows assessing three-dimensional (3D) bone structure and strength at the trabecular level (MacNeil and Boyd, 2007; Melton et al., 2007; Boutroy et al., 2008; Kazakia et al., 2008; Burghardt et al., 2010; Seeman et al., 2010; MacDonald et al., 2011). More recently, complementary methods have been proposed to computationally monitor 3D bone microstructure changes over time (time-lapse) and calculate local mechanical loading using micro-finite element (micro-FE) analysis. This has been demonstrated in mice (Schulte et al., 2013; né Betts et al., 2020; Malhotra et al., 2021) and patients (Christen et al., 2014; Mancuso and Troy, 2020) at such high spatial resolution that cellular behaviour – in the form of bone remodelling sites – can be studied and the corresponding mechanical loading can be calculated. Subsequently, these methods can be used to investigate bone’s underlying mechanoregulated remodelling process, which may be the key to the development of patient-specific therapeutic or pharmacological interventions for various bone diseases.

Typically, when investigating bone mechanoregulation under controlled experimental conditions, micro-FE models disregard subject-specific variations in external loading conditions using simplified uniaxial compressive displacement boundary conditions (SC) (Schulte et al., 2013; Mancuso and Troy, 2020; né Betts et al., 2020; Malhotra et al., 2021). However, when investigating mechanoregulation in patients, variations in day-to-day external loading are more substantial due to habitual differences and patient-specific variability in the musculoskeletal system’s performance. Distinctive tensile forces and moments are applied to joints on a routine basis to stabilise under gravitational and other external loads and create unique loading patterns (Watkins, 2009). Consequently, to investigate mechanoregulation under day-to-day loading in a personalised medicine approach, patient-specific physiological loading patterns and boundary conditions need to be estimated (Galibarov et al., 2010; Yosibash et al., 2020).

In an effort to quantify *in vivo* loading patterns using biomechanical models, several load estimation algorithms have been developed. Artificial neural network-based approaches have been proposed (Garijo et al., 2014, 2017; Mouloodi et al., 2020) but lack interpretability, which is critical for moving to diagnostic use in patients to guide local therapeutic interventions. As a result, an algebraic method introduced by Christen et al. (2012a) has been widely implemented to approximate the internal load history based on bone morphology (Christen et al., 2014; Badilatti et al., 2017; Synek et al., 2019; Cheong et al., 2020; né Betts et al., 2020). This algorithm super-imposes and scales a finite number of loading cases until a target tissue load of homogeneous strains is found. Christen and colleagues demonstrated the capabilities of such a reverse-engineering approach using an extra physiological tail-loading animal model, predicting the applied compressive loading in mouse caudal vertebra. However, remaining signal inhomogeneity remained high, ranging between 20-67%, indicating that no homogenous tissue load could be found (Christen et al., 2012). This suggests that the actual *in vivo* load distribution might differ systematically from the homogeneous assumption in humans (Christen et al., 2016; Johnson and Troy, 2018) and mice (Christen et al., 2012). By modelling homogenised strain patterns, the conventional algorithm may reduce mechanical signal inhomogeneities that have been recognised as drivers for the mechanoregulated remodelling process in bone (Frost, 1987, 2003). Thus, this model’s assumptions may not be optimal and do not fully utilise all available information in time-lapsed data of longitudinal bone imaging studies.

This study had two goals; first, to derive a validated, robust and specific method to estimate *in vivo* loading. Second, to apply this algorithm to examine *in vivo* mechanoregulation (Schulte et al., 2013) in humans and mice. We hypothesised that by extracting bone remodelling sites from time-lapsed imaging data, the relationship between bone formation in high strain regions, quiescence in medium strain regions, and resorption in low strain regions could be used in a reverse-engineering optimisation approach to determine organ-level loads. We validated our mechanoregulated approach (MR) using three unique datasets and benchmarked it with an existing load history algorithm (LH) (Christen et al., 2012). First, to calculate sensitivity, specificity and accuracy, MR and LH algorithms were applied to synthetic remodelling data derived from HR-pQCT images (Badilatti et al., 2016; Ohs et al., 2020a). Second, to test whether the algorithms are capable of predicting the loading conditions in a controlled experimental setup, both algorithms were applied to micro-CT scans of two groups of mice that had their caudal vertebra either loaded (8 N) or sham loaded (0 N) from a previous study (Scheuren et al., 2020b). Third, to assess the method’s fidelity in patients, MR and LH algorithms were applied to time-lapsed HR-pQCT scans and compared to patient-specific handgrip force measured using a dynamometer. Finally, to quantify the association between bone remodelling and mechanical stimulus, we derived a correct classification rate (CCR) (né Betts et al., 2020).

## Materials

### Human HR-pQCT images *in vivo*

HR-pQCT images (XtremeCT II, 60.7 μm voxel size, 68 kV, 1470 μA, integration time of 43 ms) were acquired from the database of a prior Innsbruck Medical University fracture study (Atkins et al., 2021). Patients gave informed consent and participated in an examination approved by the Medical University of Innsbruck ethics Committee (UN 0374344/4.31). For each patient, scans of the intact contralateral radius were taken at six time points (1, 3, 5, 13, 26, and 52 weeks) post-fracture, 9mm proximal to the endplate of the distal radius (**Figure 1**). As a functional indicator of daily mechanical load, handgrip strength was measured at 3, 6, and 12 months post-fracture using a hydraulic handgrip dynamometer. Grip strength was taken in a seated position with the elbow bent 90 degrees in flexion, measured three times and averaged. Measurements were recorded in kilograms and converted to Newtons (1 N = 9.81 Kg). Images were graded by two skilled operators using a standard visual grading score (VSG) ranging from 1 (no visible motion artefacts) to 5 (major horizontal streaks) (Whittier et al., 2020). Distal radius images of seven patients (3 males, 4 females) were included from the study by applying the following inclusion criteria. Only males or pre-menopausal female patients without fracture history of their non-dominant left distal radius were included. Only patients for whom all scans met a minimum VSG of 3 (some artefacts) and a VGS of less than or equal to 2 (very slight artefacts) in four out of the six total follow-up scans were included. The median age of the included patients was 33 years and ranged between 27 and 65 years.

**Figure 1.**
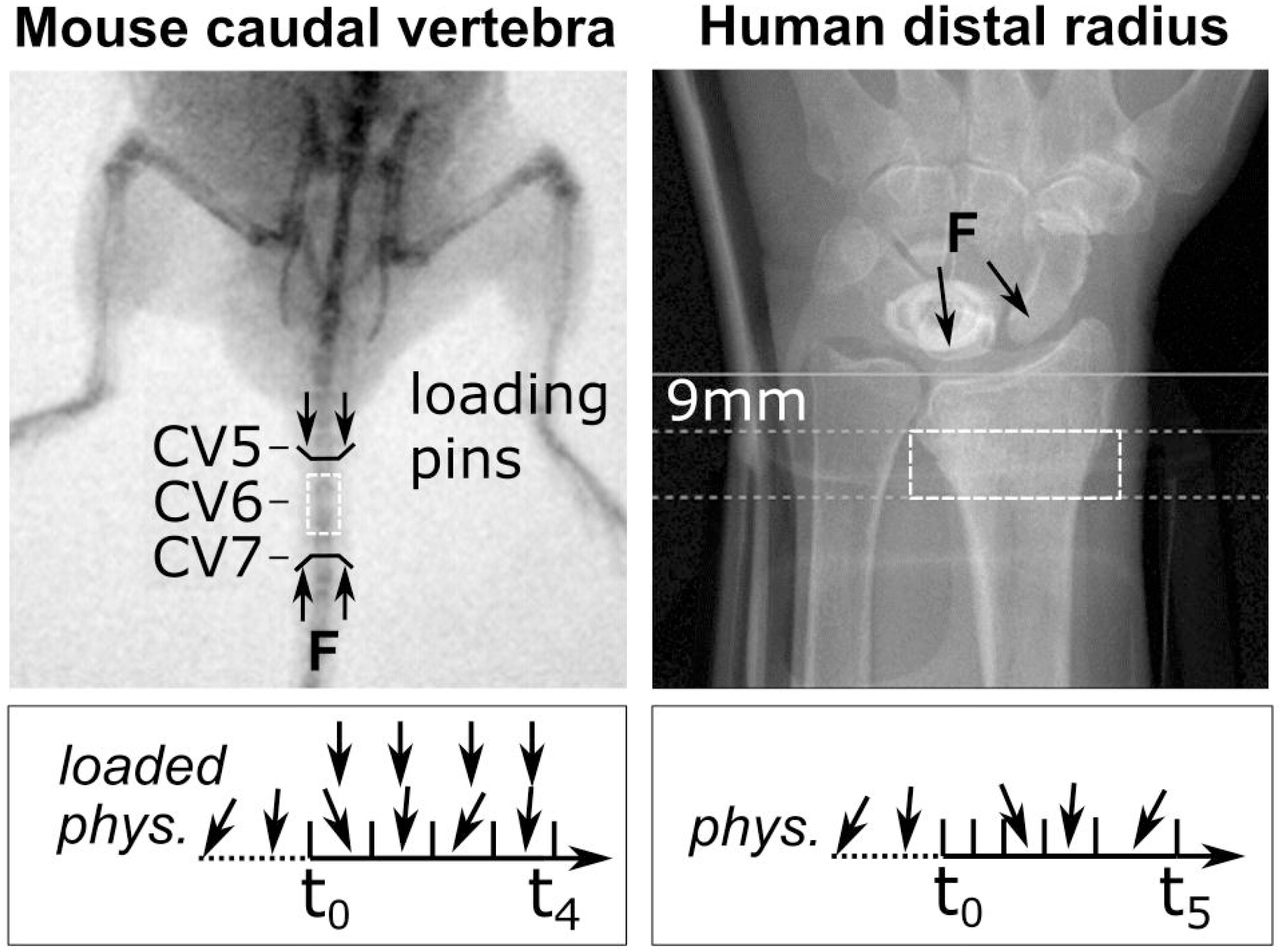
Representative fluoroscopic images of *in vivo* scanning sties. The C6 mouse caudal vertebra (dashed box, left) was scanned by micro-CT. Black lines indicate sites of loading pins in the C5 (clamped) and C7 (loaded) vertebra. A representative loading scenario is indicated below for physiologically loaded (phys.) and extraphysiologically loaded (loaded) groups throughout the study (t_0_ - t_4_). The human distal radius (dashed box, right) was scanned using HR-pQCT (Xtreme CT II). Annotations indicate the manufacturer’s recommended scanning site, 9 mm proximal to the reference line, and the arrows represent the line of action of the joint forces on the radius as a result of physiological loading. The box below indicates representative loading throughout appointments t_0_ - t_5_.

### Murine micro-CT images *in vivo*

Micro-CT images (vivaCT 40, 10.5 μm voxel size, 55 KVp, 145 μA, integration time of 350 ms, 500 projections) were acquired from a previously published mouse tail loading study (Scheuren et al., 2020b). Two groups (n = 8, each) of 11-week old female C57BL/6J strain mice were scanned at the 6th caudal vertebrae (CV6) at weekly intervals for five weeks. The sixth caudal vertebra of the animals in the loaded group was subject to mechanical loading through stainless steel pins inserted into the adjacent vertebrae (**Figure 1**). Compressive loading was applied three times per week for five minutes at 10 Hz and 8 N. Animals in the control group were subject to sham loading (0 N) (see Scheuren et al., 2020b).

## Methods

### Image processing

After rigid image registration (Schulte et al., 2014), distal radius images were upscaled to 30.5 μm (Ohs et al., 2020a), and caudal vertebra images were kept at 10.5 μm native resolution. Images were Gauss filtered to reduce noise (sigma 1.2, support 1). Human distal radius and mouse vertebra scans were binarised using a threshold of 320 mg/cm^3^ and 580 mg/cm^3^, respectively (Hosseini et al., 2017; Scheuren et al., 2020a). Trabecular regions were automatically contoured from binarised images. For the human distal radius images, an approach described by Ohs et al. (2020b) was used; for the mouse vertebra images, a method described by Kohler et al. (2007) was used. Finite element meshes were generated by converting all voxels to 8 node hexahedral elements and assigning a Poisson’s ratio of 0.3 as well as Young’s modulus of 6.8 GPa for the human distal radius (Christen et al., 2013) and 14.8 GPa (Webster et al., 2008) for the mouse vertebra. Remaining interior voxels located within the bone cavity were assigned a value of 2 MPa and a Poisson’s ratio of 0.3 (Webster et al., 2008). For the mouse caudal vertebra, intervertebral discs with a Young’s modulus of 14.8 GPa were approximated and added to the proximal and distal ends of the vertebra (Webster et al., 2008; Schulte et al., 2013).

### Micro-finite element analysis

Axial and shear forces were applied to the target tissue’s distal and proximal surfaces using a 1% displacement boundary condition. Torsion and bending moments were applied, centred around their corresponding axis, with a 1° displacement. Six uniaxial loading directions were defined: compressive force in the axial direction (C), lateral shear force in the (SX), dorsal shear force (SY), axial moment around the long axis (MZ), lateral bending moment (BX) and dorsal bending moment (BY). Linear FE calculations were carried out using ParOsol (Flaig and Arbenz, 2011); the solver was run at the Swiss National Supercomputing Centre (CSCS, Lugano, Switzerland) using 128 CPUs. Strain energy density (SED) was used as a mechanical signal for bone remodelling. Unit load cases were derived by rescaling applied force magnitudes to 1 N, moment magnitudes to 1 Nmm, and resulting SED distributions accordingly (Christen et al., 2012). Three multiaxial loads were defined using a method of scaling and superimposing unit load cases modelling the aggregated effect of physiological load over time: combined compression and shear (CS = 0.5 C + 0.25 SX + 0.25 SY), combined compression and bending (CB = 0.5 C + 0.25 BX + 0.25 BY) and a combined 6 degree of freedom load (6DoF) with equal proportions of load in all six uniaxial directions.

### Remodelling algorithm

Five patients (4 females, 1 male) with VGS lower than 2 were randomly selected from the initial patient cohort due to the high computational cost of the remodelling simulation. Geometries derived from baseline HR-pQCT scans were adapted towards previously defined uniaxial (C, SX, SY, MZ, BX, BY) and multiaxial (CS, CB, 6DoF) loads using a modified advection-based remodelling algorithm (Badilatti et al., 2016; Ohs et al., 2020a). In short, a regularised density, that matched binary BV/TV while preserving greyscale value on the bone surface, was converted to Young’s modulus using a linear relationship (Mulder et al., 2007) and used as input for the remodelling algorithm (Ohs et al., 2020a). The advection-based remodelling process, as described in **Figure 2**, was limited to the trabecular region and performed for each of the nine *in silico* loading experiments for 40 remodelling steps. SED and applied force magnitudes derived from micro-FE analyses were rescaled to target sample-specific homeostatic remodelling with comparable amounts (<2% difference) of bone formation and resorption. Five images were subsampled for each patient and loading experiment using a Pearson correlation coefficient with a threshold of 0.95 to model HR-pQCT follow-up scans’ temporal resolution.

**Figure 2.**
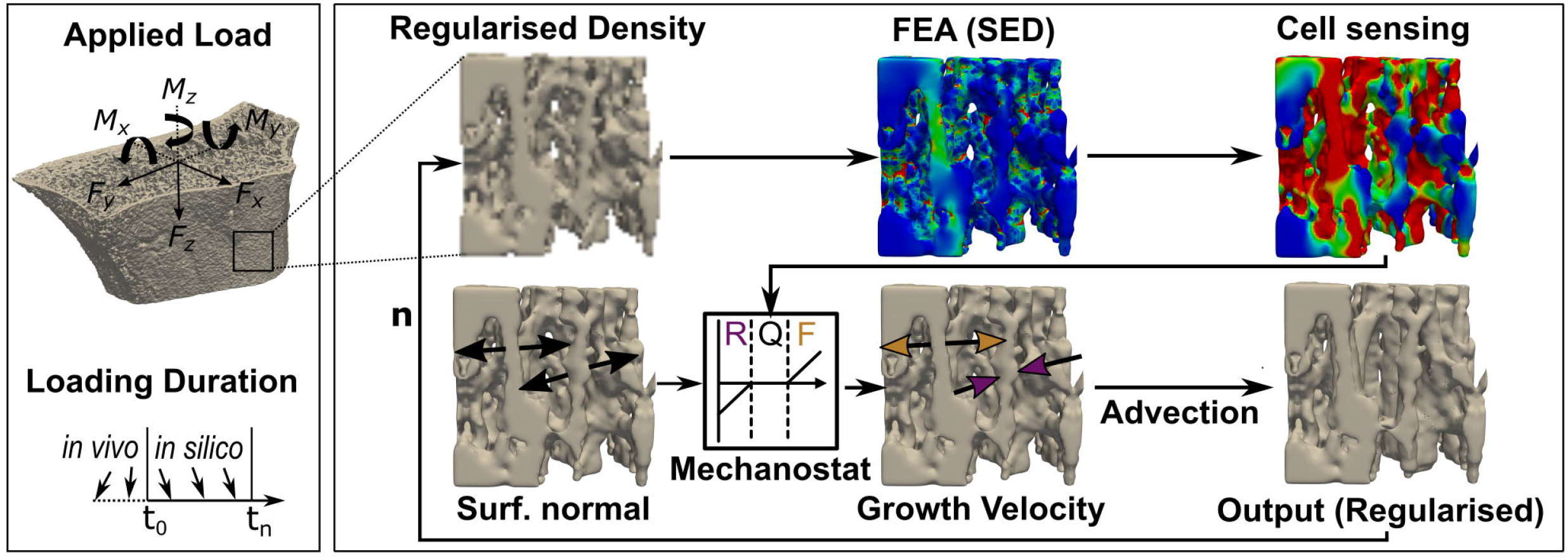
Schematic workflow to derive bone geometries from advection-based remodelling simulations. Input, greyscale HR-pQCT images of the distal radius were first Gauss-filtered and regularised before finite element modelling. Strain energy density (SED) was derived from a linear finite element analysis (FEA), and cell sensing was mimicked through mechanical signal dilation with a fixed radius of 50 μm. Tissue was remodelled using a SED dependent velocity of ±8000 μm/year/MPa and a maximum velocity of ±12 μm/month in regions where SED exceeded or fell short of the average tissue load (0.02 MPa) by ±2%, and the growth direction was simulated normal to the bone surface. An advection step performed the surface movement, either resorption (R, purple) or formation (F, yellow), and a remodelled output regularised image was derived. Quiescence (Q) was modelled as no surface movement. This process repeats with the regularised output image as input for the next iteration (n).

### Mechanoregulation-based load estimation

The mechanoregulation-based load estimation (MR) was performed in two steps and followed established mechanoregulation principles (Wolff, 1892). The algorithm operated on the bone surface S(x), which was defined as the interface between the bone and the background using a 3D von Neumann neighbourhood with a radius of 1 voxel. New bone was presumed to be formed in high mechanical signal regions, quiescent in regions of medium mechanical signal and resorbed in regions of low mechanical signal (see **Figure 3**). Regions of formation RV_f_, quiescence RV_q_ and resorption RV_r_ were calculated by overlaying two subsequent binary images aligned using rigid registration. Each surface voxel was assigned a rank rg_RS_ according to its remodelling event (resorption = 1, quiescence = 2, formation = 3). Accordingly, an ordinal definition of the mechanical signal rg_SED_ was specified with increasing rank for increasing signal magnitude. The monotonic relationship between rg_RS_ and rg_SED_ represents a mechanoregulated behaviour between surface remodelling events and mechanical signal.

**Figure 3.**
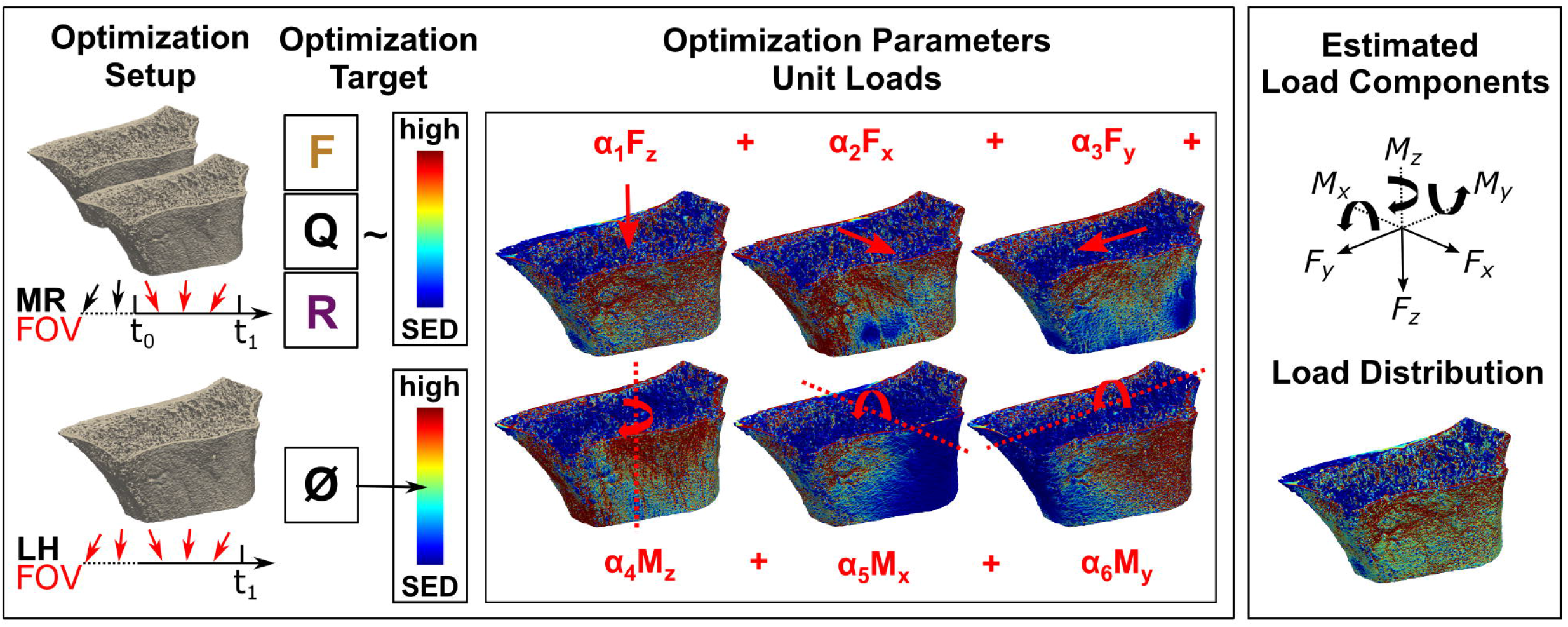
Overview of the mechanoregulation-based load estimation algorithm (MR) and morphology-based load history algorithm (LH). (Top left) *In vivo* loading is assessed by MR between two consecutive images, outlining the algorithms field of view (FOV). By overlaying registered, longitudinal images, remodelling regions are identified to find a loading scenario maximising the correlation between formation (F) in regions of high strain, quiescence (Q) in regions of medium strain and resorption (R) in regions of low strain. (Bottom left) In comparison, LH estimates the complete *in vivo* load history with no option to limit its FOV and targets a homogeneous strain distribution of medium strain (0.02 MPa). (Center) For the optimisation in both algorithms, micro-FE models are created covering all physiologically possible loading directions. During the optimisation unit loads are scaled until the optimisation target is achieved, providing (Top right) individual load components (i.e. forces and moments) as well as (Bottom right) a combined load distribution.

In the first optimisation step, Spearman’s rank-order correlation between rg_RS_ and rg_SED_ was maximised by scaling a set of previously defined unit load cases U_(i,unit)_(x) with load composition factors c_i_ (with c_i_ ∈ [0, 1]). Where U_(i,unit)_(x) is the SED distribution due to unit load i on the bone surface S(x). The superimposed unit loads defined a potential compounded mechanical stimulus with known unit load proportions within each iteration. A gradient-free Nelder-Mead method with a tolerance of 10^−4^ was used to optimise the following resulting equivalent minimisation objective function r.

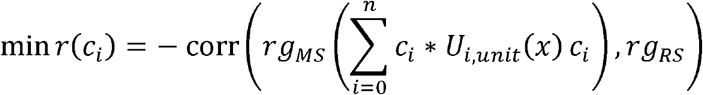

The resulting load composition c_i_ determined the best combination of unit loads to associate bone formation in regions of high, quiescence in areas of medium, and resorption in regions of low signal. However, no assumptions on the magnitude of the mechanical signal were made. To derive the final mechanical load, a second optimisation procedure matching the compounded signal with bone’s overall remodelling response was performed. Bone formation rate (BFR), bone resorption rate (BRR) and net remodelling response (NRR = BFR - BRR) were calculated from the registered binary images (Lambers et al., 2011; Schulte et al., 2011). To calculate NRR_SED_ as predicted by the mechanical signal, we defined a ternary classifier function f_j_ considering two thresholds for sites of formation T_f_ and sites of resorption T_r_ according to Tourolle né Betts et al. (2020). The thresholds T_f_ = 0.0204 MPa and T_r_ = 0.0196 MPa were chosen based on average bone loading values of 0.02 MPa from previous studies (Christen et al., 2012, 2013). To observe both, formation and resorption, in the simulations a narrow 4%-wide lazy zone was implemented. At each iteration NRR_SED_ was calculated by scaling the compounded mechanical signal using a second scaling factor r, and the prediction of the classifier function f_j_(r ∗ ∑c_i_ ∗ U_i,unit_(x)) within was used in the current study. A gradient-free Nelder-Mead method with a tolerance of 10^−4^ was used to minimise the difference between NRR_SED_ and NRR_GT_ using the following objective function k(r).

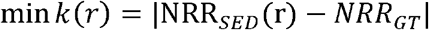

For consistency with Christen et al., (2013), scaling factors c_i_ and r were incorporated into a single scaling factor s_i_ = r ∗ c_i_, which combines magnitude and number of load cycles applied over time. Assuming each load case acted equally long over time and was applied sequentially, loading magnitude α_i_ was calculated as α_i_ = √(1/6 ∗ s_i_) for the six applied unit load cases.

### Morphology-based load history estimation

Following a previously published approach (Christen et al., 2013), we implemented a load history algorithm (LH). Unit load cases were scaled using load composition factors s_i_ until the most homogeneous distribution is found (k = 0.02 MPa) (see **Figure 3**). Scaling factors s_i_ were calculated using a non-negative linear least-squares optimisation technique and load magnitudes α_i_ were calculated as previously described. Further, a calibrated version of LH was implemented (cal. LH). In its native implementation, LH evaluates the load history before the imaging timepoint. In longitudinal studies, physiological loading during the study may change compared to loading before the study. To reduce this initial bias from prior loading, the scaling factors estimated by LH α_i,t-1_ from the previous, baseline image were subtracted from the estimated scaling factors α_i,t_ of the current timestep.

### Multiclass receiver operator characteristic (ROC)

A multiclass ROC averaging approach was used to assess the accuracy of the *in silico* load estimation. The Euclidean distances between the estimated and all possibly applied load vectors were calculated as a scalar error quantification. The multiclass prediction of all 9 *in silico* loading scenarios was reduced to multiple sets of binary predictions (true, false) for each scenario. A true positive rate (TPR) was assessed over a false positive rate (FPR) at different thresholds, and the area under the curve (AUC) was calculated. Following Mandrekar (2010), AUC of 0.5 suggested no discrimination, 0.7 to 0.8 was considered acceptable, 0.8 to 0.9 was deemed excellent, and larger than 0.9 was considered outstanding. The receiver operator characteristic (ROC) was calculated for each scenario, and the results were averaged to calculate a macro average (mac). Further, a prevalence-weighted micro average (mic) was calculated treating data as an aggregated result. These averages describe the overall performance of the multiclass classification (Asch, 2013).

### Tissue-level error quantification

Forces and Moments act on different scales and are not directly comparable in magnitude. However, their resulting strain distributions may help understand their impact on tissue scale. In contrast to the *in silico* data, in the *in vivo* data, no ground truth was available to validate the results directly. For the animal data, an anticipated SED distribution was derived based on the experimental assumptions in order to conditionally validate the MR algorthim. The loaded group was subject to an 8 N cyclic load; consequently, a local reference SED distribution was derived for an 8 N load for the loaded group. In accordance with Christen et al. (2012a), a 4 N compressive load was assumed for unloaded animals and the according to SED distribution was derived. The error between LH and MR’s load distributions to the reference distributions was calculated for each voxel by subtracting the target’s estimated distribution. Voxels were binned according to derived remodelling regions, resulting in error distributions for areas of formation, resorption and quiescence.

### Conditional probabilities of remodelling events and correct classification rate

Conditional probability (CP) curves were calculated for the previously identified remodelling events on the bone surface, in accordance to Schulte et al. (2013a), to connect the mechanical environment (SED) as estimated by the algorithms with remodelling sites. Load distribution, resulting from the estimated loads, was normalised using the 99^th^ percentile and binned at 1% steps for each remodelling event. A group-wise and bin-wise normalisation were used to calculate CP curves for each data set (Schulte et al., 2013). A correct classification rate (CCR) adapted from Tourolle né Betts et al. 2020 was calculated to summarise mechanoregulation. This correct classification rate measures the fraction of correctly identified remodelling events using the conditional probability curves.

### Statistics

Statistical analysis was performed using Python 3.8.0, NumPy 1.19.2 and SciPy 1.5.3. Data were tested using an omnibus test of normality based on D’Agostrino and Pearson that combines skew and kurtosis (d’Agostino, 1971; D’Agostino and Pearson, 1973). Non-normal parameters were presented as median ± 95% confidence interval (CI) and compared using nonparametric tests: the Wilcoxon-Mann-Whitney test was used for independent and the Wilcoxon-Signed-Ranks test was used for matched samples. To measure the association between MR and LH predictions and their correlation with grip strength, linear regression analysis was performed; for non-normal parameters, Spearman’s rank-order correlation cofficients were computed to assess the relationship between variables. Normal parameters were presented as mean ± 95% CI and compared using parametric methods: the Student’s t-test was used for independent samples and a paired T-test was used for matched samples. For linear regression analysis of normal parameters, Pearson product-moment correlation coefficients were computed. Holm-Bonferroni correction was used for multiple comparisons to reduce the possibility of a type I error. For all tests, a p-value smaller 0.05 was regarded as statistically significant. Otherwise, significance levels are reported.

## Results

### Generation of adapted bone geometries *in silico*

For the *in silico* experiments, the goal was to generate adapted bone geometries with constant remodelling rates and known mechanical loads to benchmark the algorithms. The *in silico* applied force magnitudes were varied until homeostatic remodelling was achieved, resulting in forces between 100 N and 600 N. Average BV/TV of the baseline trabecular geometries was 0.12 ± 0.06 and increased to 0.13 ± 0.06 at step 40. An initial drop in BV/TV was observed within the first eight steps of the simulation’s initialisation period and was excluded from further analysis. The temporal resolution between the resulting advection steps needed to be reduced to achieve physiological and constant remodelling rates comparable to *in vivo* follow up periods. Linear regression analysis showed a significant negative correlation (R^2^ = 0.97, p < 0.01) between remodelling rates and Pearson’s R between two subsequent images. Hence, Pearson’s R was regarded as a reliable subsampling criterium. Timepoints were included when a threshold of 0.95 was reached between images resulting in six to eight scans for each geometry and loading scenario. The last six subsampled timepoints for each experiment and patient were selected for further analysis. This procedure provided highly controlled remodelling rates of 13.79 ± 0.13% between scans.

### Sensitivity, specificity and accuracy *in silico*

A multiclass ROC analysis was used to assess Sensitivity, Specificity and Accuracy. Average AUCs were high for MR calculated using micro (AUC = 0.98) and macro (AUC = 0.97) averaging. This high value was due to an outstanding performance when classifying uniaxial loads (AUC = 1) (**Table 1**) and dropped for multiaxial loading cases (AUC = 0.91). An overshadowing of the shear component by compression was observed for CS, resulting in a considerable AUC drop (see **Table 1**). Still, MR exceeded the performance of LH in all categories (**Figure 4**). Overall, LH only resulted in acceptable micro (AUC = 0.61) and macro (AUC = 0.73) averages, and a below random prediction (AUC = 0.45) was observed for the 6DoF load case. Overall, AUC improved for the calibrated implementation for macro (AUC = 0.79) and micro (AUC = 0.71) averages; however, it was not consistently higher in all categories. At the optimal ROC cut point, load configurations were correctly identified with a high sensitivity of MR. Additionally, the ratio of correctly identified mismatches manifested in high specificity, resulting in an outstanding overall accuracy of MR (**Figure 4**, upper left panel). In comparison, sensitivity, specificity and accuracy of LH were significantly lower (p < 0.01), yielding only an acceptable differentiation between the applied loading. The calibrated implementation of LH did not achieve significantly higher accuracy compared to the native LH approach and was not further investigated.

**Figure 4.**
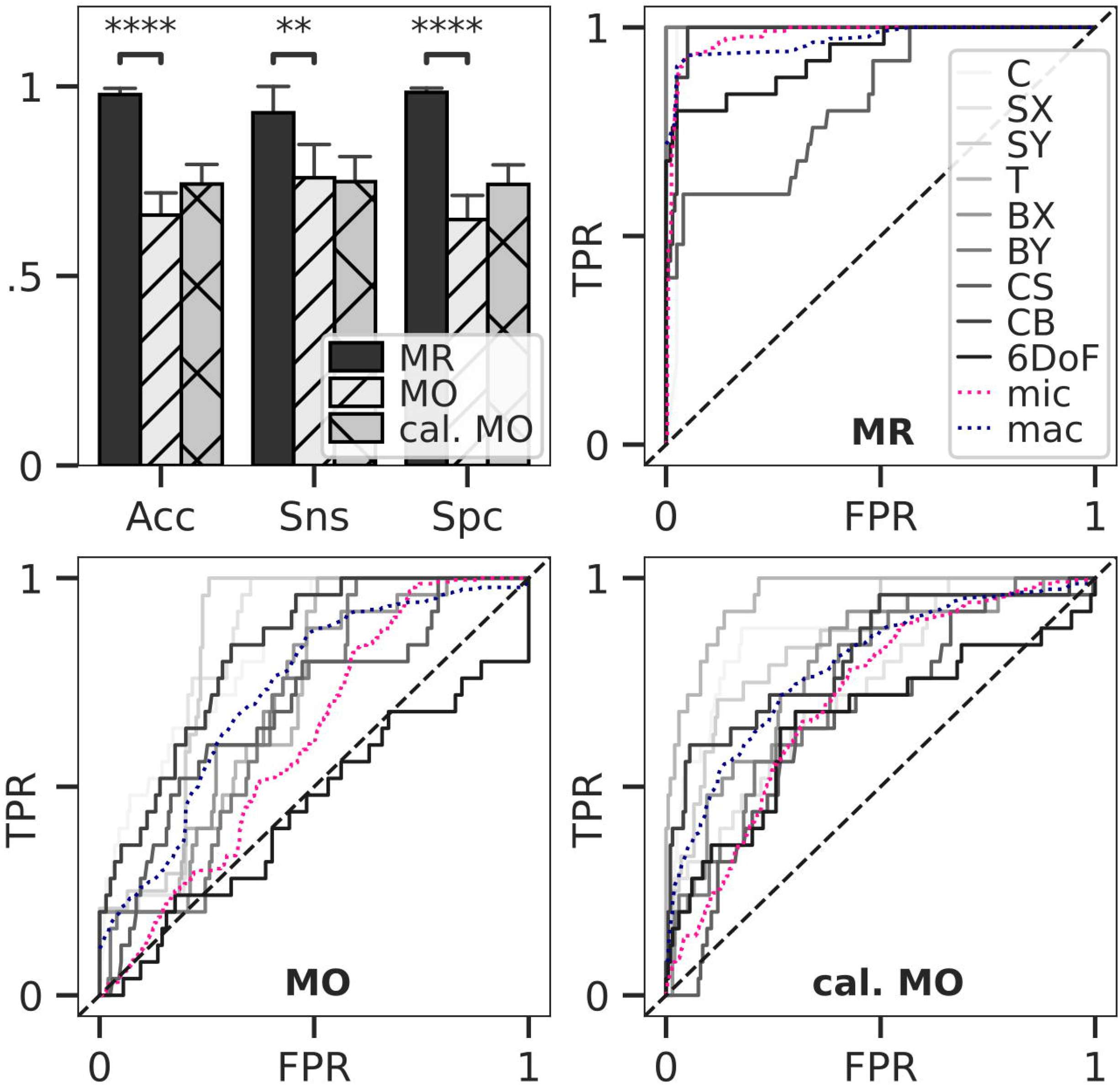
Classification accuracy and ROC for load estimation. Loads of *in silico* adapted bone geometries (n=5) with nine different loading boundary conditions were estimated and compared to the simulated target load serving as ground truth. Accuracy, Sensitivity and Specificity for estimated-optimal thresholds were calculated (upper left). Bars show mean, and error bars 95% confidence interval. All differences between means with p < 0.05 are indicated (**, p < 0.01; ****, p < 0.0001; two-tailed paired t-test). Thresholds were derived from multiclass ROC for MR (upper right), LH (lower left) and calibrated LH (lower right)

**Table 1.** ROC derived area under the curve (AUC) for MR, LH and calibrated LH for uniaxial loading cases and multiaxial loading. Averages were calculated based on aggregated averaging (macro) and a prevalence-weighted average (micro).

|  | Uniaxial |  |  |  |  |  | Multiaxial |  |  | Average |  |
| --- | --- | --- | --- | --- | --- | --- | --- | --- | --- | --- | --- |
|  | C | SX | SY | T | BX | BY | CS | CB | 6dof | Micro | Macro |
| MR | 0.98 | 1 | 1 | 1 | 1 | 1 | 0.82 | 0.98 | 0.93 | 0.98 | 0.97 |
| LH | 0.83 | 0.81 | 0.83 | 0.71 | 0.7 | 0.68 | 0.69 | 0.82 | 0.45 | 0.61 | 0.73 |
| Cal.<br>LH | 0.87 | 0.74 | 0.85 | 0.96 | 0.78 | 0.75 | 0.66 | 0.82 | 0.66 | 0.71 | 0.79 |

### Association between different load estimation algorithms *in silico*

Linear regression between MR and the target load resulted in α_target_ = 1.28 ∗ α_MR_ + 2.64 (R = 0.83, p < 0.05), slightly underestimating loading magnitude. In comparison, LH showed a weaker correlation and overestimated loads (α_target_ = 0.86 ∗ α_MO_ - 1.80, R = 0.45, p < 0.05). The calibrated version of LH showed a slightly higher correlation; however, loading magnitudes were underestimated by orders of magnitude indicating that the calibrated version of LH should only be used in combination with a valid calibration equation (α_target_ = 8.36 ∗ α_calMO_ + 36.18, R = 0.5, p < 0.05).

### Subject-specific load in the mouse caudal vertebra *in vivo*

One animal of the loaded group was excluded from the analysis due to convergence issues during the finite element analysis. The axial compressive force was predicted as the most dominant loading component for all time pionts using MR (loaded 6.11 ± 1.15 N, control 4.40 ± 1.37 N). Estimations in the loaded group were consistently higher compared to the unloaded group (3.73 ± 2.13 N) reaching significantly (p < 0.05) higher levels after two weeks (see **Figure 5a**). In comparison, the estimations of the axial compressive force by LH only reflected the experimental conditions in the loaded group after the 2-week time point, predicting 5.24 ± 1.42 N in control and 6.40 ± 3.72 N in the loaded group. Using MR, a non-negligible M_x_ moment was predicted in both the loaded (3.97 ± 4.00 Nmm) and control (3.17 ± 1.03 Nmm) groups. Notably high bending moments (> 4 Nmm) in the loaded group were only observed for individual mice, causing large confidence intervals in the predicted M_x_ component of the loaded group. In comparison, M_z_ was the largest moment load component indicated by LH for loaded (13.41 ± 0.51 Nmm) and control (14.97 ± 0.33 Nmm) groups, and was significantly (p < 0.05) higher compared to M_z_ indicated by MR in the loaded (1.41 ± 0.58 Nmm) and control (4.56 ± 1.04 Nmm) groups. Errors for loading estimated by MR were normally distributed (see **Figure 5b**). In comparison, errors for loading estimated by LH were skewed left in regions of resorption resulting in a systematic overestimation of strain in these areas (see **Figure 5c**). Additionally, mean absolute error was significantly (p < 0.01) smaller for estimations by MR (f: 0.0051 ± 10^−5^ MPa, q: 0.0057 ± 10^−5^ MPa, r: 0.0042 ± 10^−5^ MPa) compared to LH (f: 0.0071 ± 10^−5^ MPa, q: 0.0070 ± 10^−5^ MPa, r: 0.0081 ± 10^−5^ MPa).

**Figure 5.**
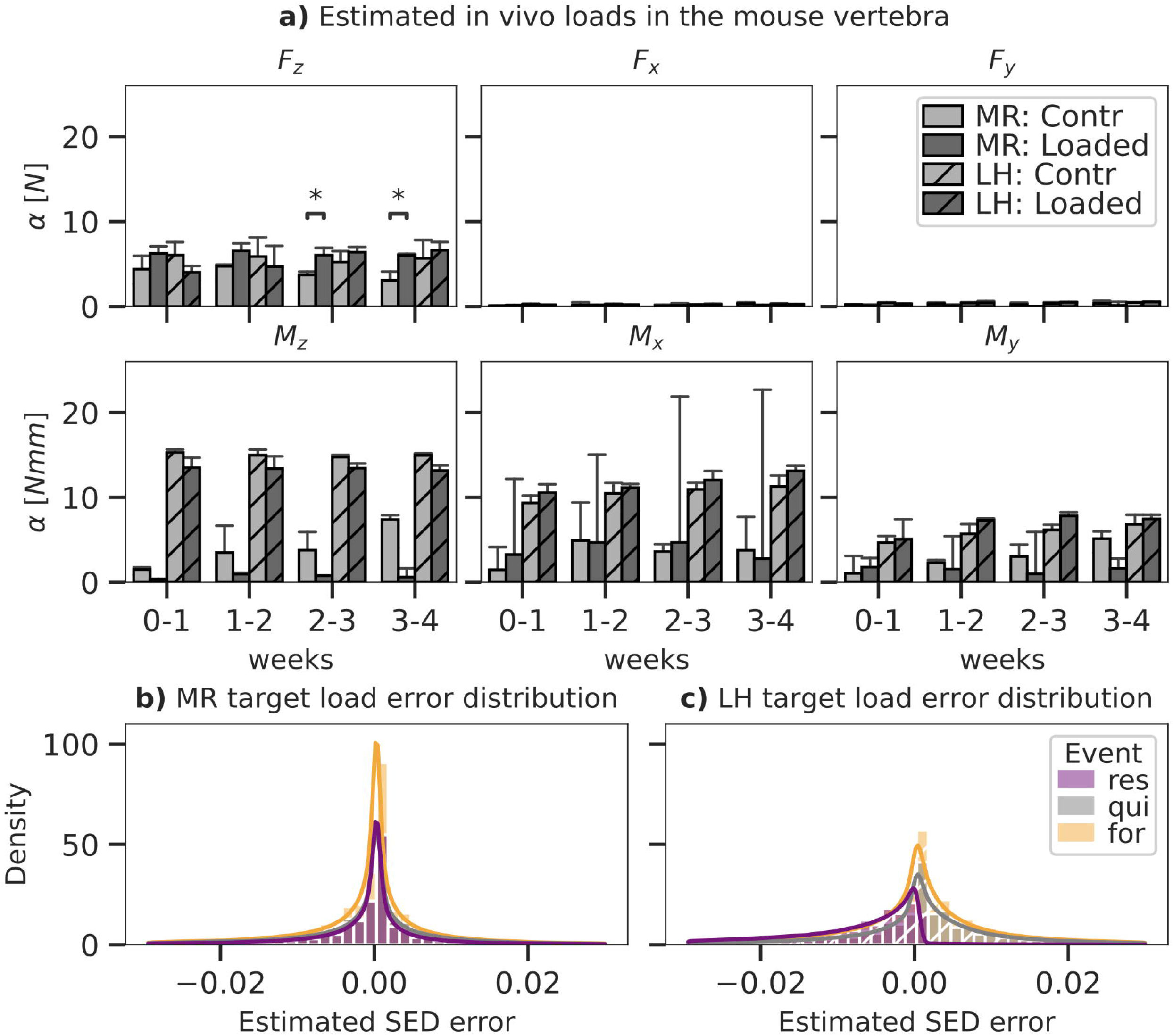
Load components and error as predicted by MR (solid) and LH (dashed) for mouse caudal vertebra (n = 8) subjected to physiological (Contr) and extra physiological loading (Loaded). Bar plots in panel (a) show mean predicted load and standard error (SE) for each component of a six-degree freedom load. Significant differences in prediction between MR and LH with p < 0.05 are indicated (*, p < 0.05; Mann-Whitney-Wilcoxon, Bonferroni). By MR and LH predicted strain energy density (SED) distributions were compared to an anticipated target load case and distribution derived from the experimental conditions (contr: 4 N in Fz, loaded: 8 N in Fz). Local error distribution was assessed between estimated and target SED for MR **(b)** and LH **(c)** and grouped in regions of formation, resorption and quiescence, as derived from time-lapsed micro-CT images.

### Patient-specific load in the human distal radius *in vivo*

Compressive force, F_z_, was the largest loading component compared to the other unit load cases in the distal radii using both MR (F_z_ = 0.43 ± 0.33 kN) and LH (F_z_ = 0.42 ± 0.27 kN); however, F_z_ did not reach a significantly higher magnitude than F_x_ (0.14 ± 0.09 kN) or F_y_ (0.28 ± 0.13 kN) (see **Figure 6a**). This may be attributed to the large variations in F_z_ predicted by MR and LH across subjects. Mean estimated F_z_ was in good agreement between LH and MR. Using MR, estimated loading was consistent over the 12 month interval showing no significant difference between time points. Loads estimated using MR showed more considerable variation than LH, which may be due to registration artefacts or variations in image quality between timepoints.

**Figure 6.**
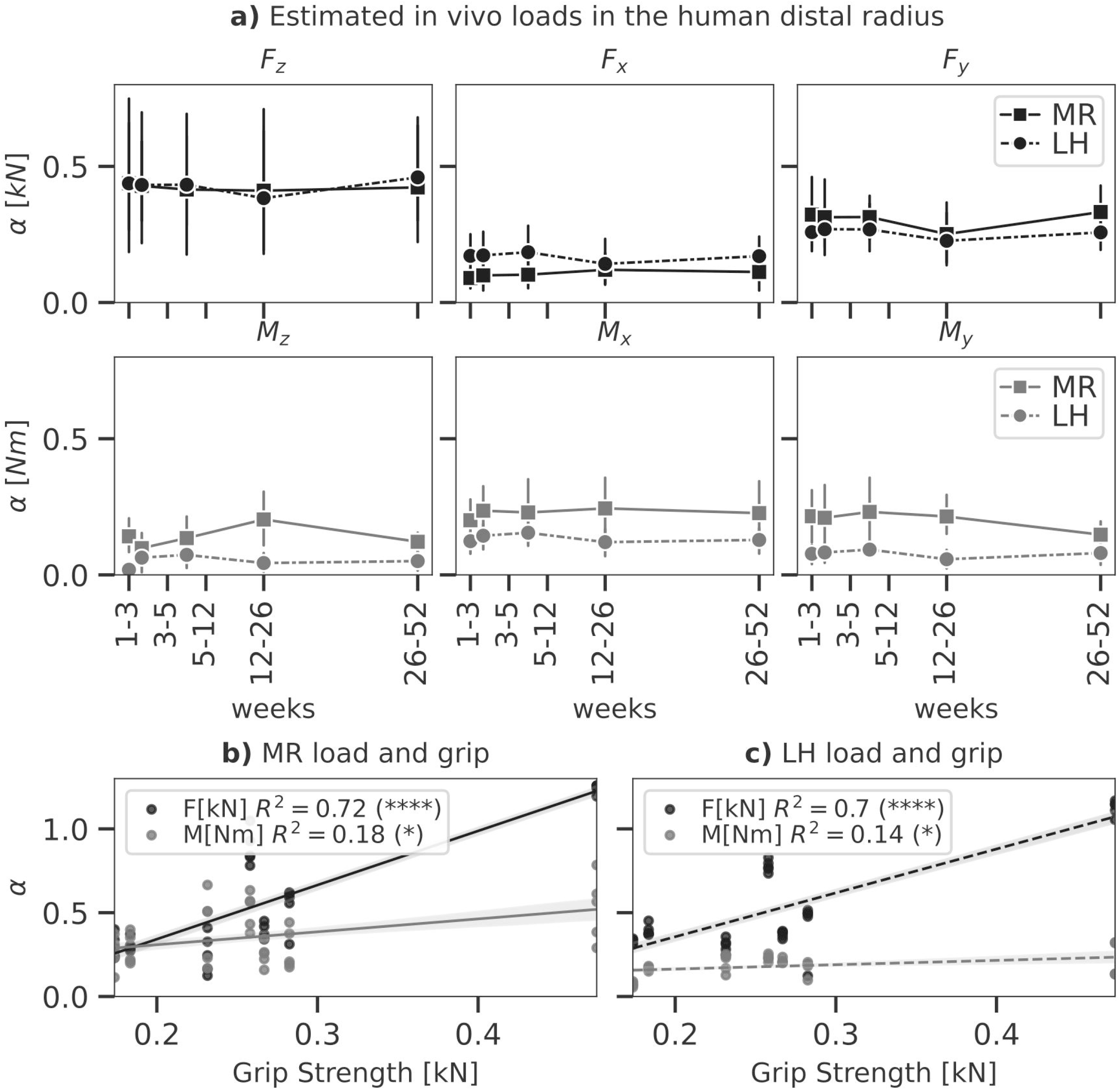
Load as predicted by MR (solid) and LH (dashed) of physiological load in the human distal radius (n = 7). In panel (a), line plots show mean predicted load and 95% confidence intervals for each component of a six degree of freedom load. No significant differences were found between MR and LH (p < 0.05, paired t-test). Linear regressions between grip strength and total force ∑F_i_ and moment ∑M_i_ as predicted by MR (b) and LH (c) were calculated. Significant correlations indicated (*, p < 0.05; ****, p < 0.0001).

Grip strength of individuals was assessed to investigate these variations in compressive force between subjects. Simple linear regression was calculated to predict loads estimated by MR (moment M in Nm and force F in kN) based on grip strength G in kN (see **Figure 6b**). For F, a significant regression equation (F(G) = 3.22 ∗ G - 0.30) was found (p < 0.01) with R^2^ = 0.72. This correlation between grip strength and forces in the distal radius has been found before and may explain variations among subjects as F increased 3.22 kN for each kN of grip strength. For M, a significant regression equation (M(G) = 0.77 ∗ G + 0.15) was found (p = 0.01) with R^2^ = 0.18. As such, moment load in the distal radius was less associated with grip strength compared to forces. Simple linear regression for loads estimated by LH reflected a similar trend with a slightly weaker association (see **Figure 6c**). For F, a significant regression equation (F(G) = 2.61 ∗ G - 0.16) was found (p < 0.01) with R^2^ = 0.70. For M, a significant regression equation (M(G) = 0.26 ∗ G + 0.11) was found (p = 0.03) with R^2^ = 0.14.

### Local mechanoregulation *in silico* and *in vivo*

Mechanoregulation analysis of MR and LH was conducted and compared to the results of a commonly used simple compression finite element analysis (SC). SED distributions were normalised using the 99^th^ percentile resulting in median normalisation values of 0.071 ± 0.06 MPa for MR, 0.04 ± 0.01 MPa for LH, and 5.28 ∗ 10^−7^ ± 0.01 MPa for SC. Mechanoregulation curves (**Figure 7a**) showed systematic bone remodelling behaviour where bone was most likely to be formed in high SED regions, quiescent in medium SED areas and resorbed in regions of low SED as visually indicated in (**Figure 8)**. The *in silico* model’s purely mechanically driven gaussian process was only fully recovered using MR. This anticipated distribution can be seen in the lower-left panel of **Figure 7a**, showing models generated and analysed using the same SC boundary condition. In comparison, LH’s cp indicated an unphysiological change in curvature localised just above 50% strain.

**Figure 7.**
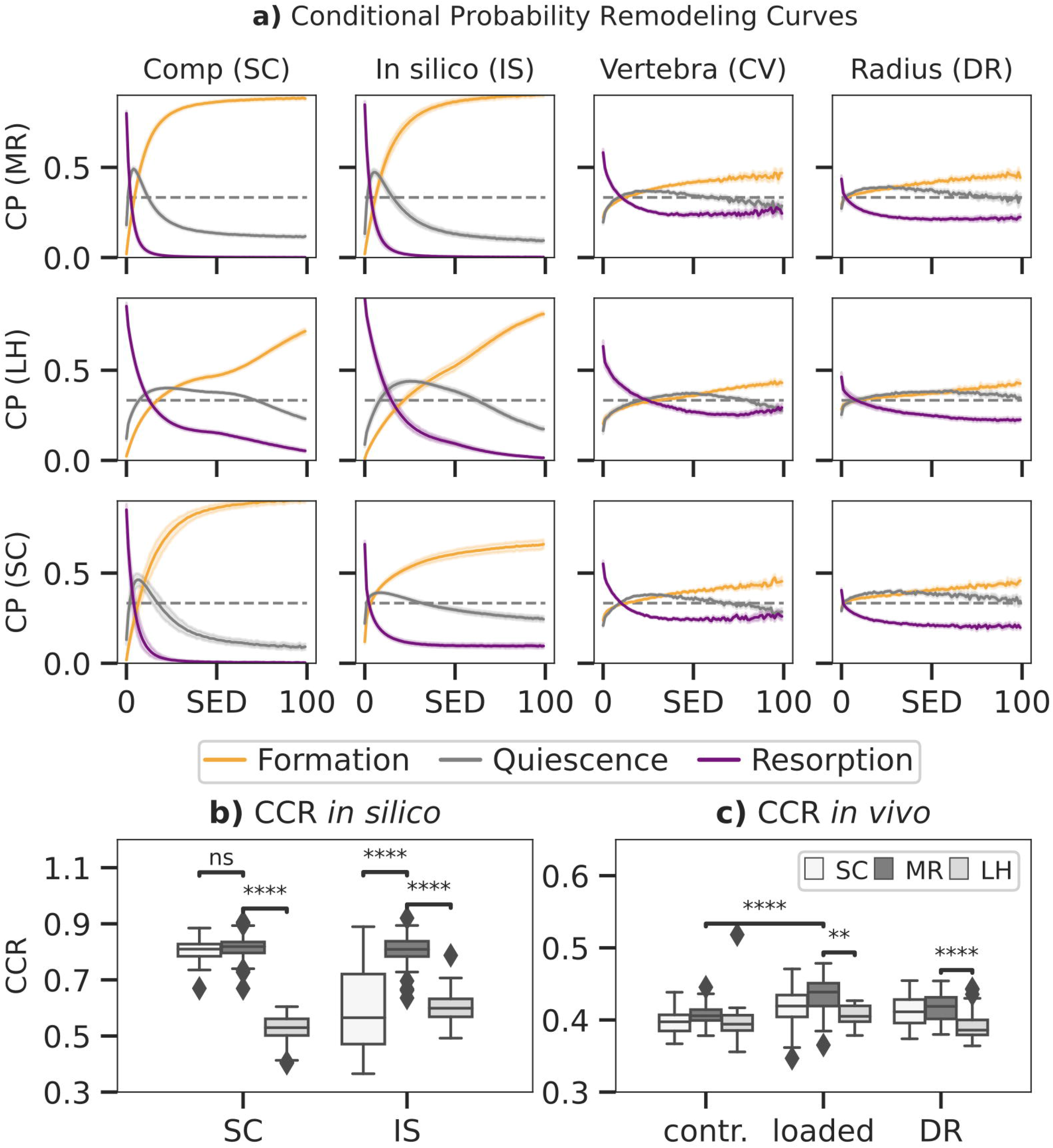
Conditional remodelling probabilities (CP) connecting the mechanical environment (SED) as estimated by MR, LH and simple compression (SC) with remodelling sites for simple compression (SC), in silico loading (IS), i*n vivo* vertebra (CV) and distal radius (DR) datasets. Normalised SED distributions were used to calculate the conditional probability **(a)** for events of formation, quiescence and resorption to occur at distinct strain levels. Remodelling sites as predicted by the estimated SED were compared to the ground truth, and a correct classification rate (CCR) for *in silico* data **(b)** and *in vivo* data **(c)** was calculated. Boxplots indicate the median and interquartile range. Observations outside the 9-91 scope plotted as outliers. Differences in prediction within and between groups with p < 0.05 are indicated (***, p < 0.001; ****, p < 0.0001; two-tailed paired t-test within groups, two-tailed individual t-test between groups).

**Figure 8.**
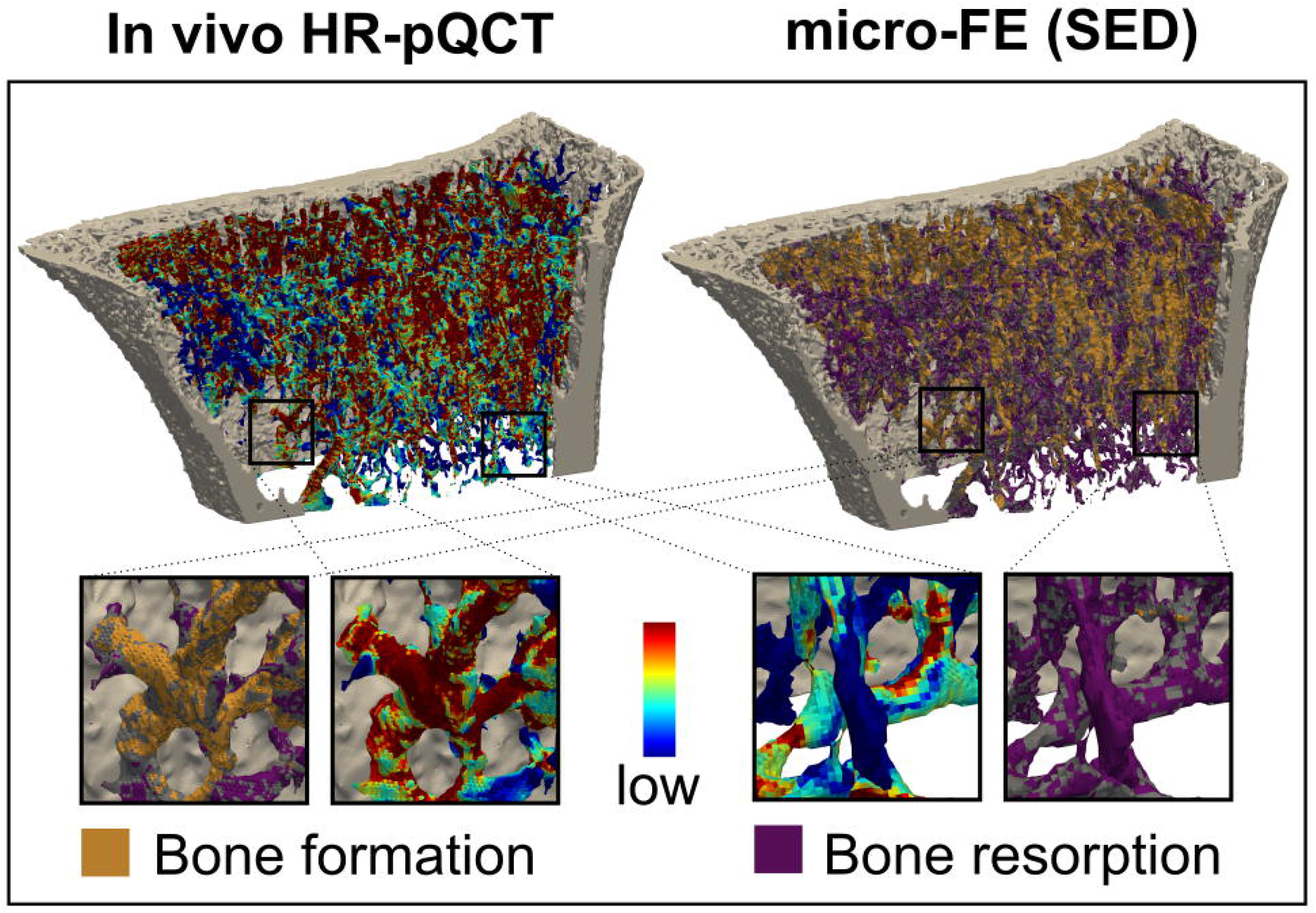
Comparison of remodelling sites with the mechanical environment. Longitudinal *in vivo* HR-pQCT scans identified bone formation, quiescence and resorption and were directly compared to the local mechanical environment. The inset shows an enlarged view of the correspondence between bone formation and high signal and low signal resorption.

To quantify the overall remodelling behaviour, CCR was calculated, measuring correctly classified remodelling events. CCR was significantly higher in the *in silico* data (**Figure 7b)** than *in vivo* data (p < 0.01) as seen in **Figure 7c**. For the *in silico* data, MR achieved significantly higher CCR (CCR = 0.81) compared to LH (CCR = 0.55, p < 0.01). Comparison between the SC (CCR = 0.80) benchmark and MR (CCR = 0.81) showed no significant differences, demonstrating high *in silico* performance of MR. Within the *in vivo* mouse data, no significant differences in CCR were found in the unloaded group (CCR = 0.40) between approaches. However, in the loaded group, CCR predicted using MR (CCR = 0.43) was significantly higher compared to LH (CCR = 0.40, p < 0.01) and significantly higher compared to the unloaded group. Finally, within the human distal radius, significantly larger association was found between strain derived from MR (CCR = 0.42) compared to LH (CCR = 0.38, p < 0.01), and a higher trend compared to SC (CCR = 0.41).

## Discussion

With the increasing prevalence of bone mechanoregulation studies, this work aimed to extend a previously developed load estimation algorithm (LH) (Christen et al., 2012) by allowing for tissue strain inhomogeneities in our mechanoregulated load estimation approach (MR). These localised differences in mechanical signal may drive bone’s remodelling response and help understand bone mechanoregulation. We provided validation for both algorithms using *in silico* generated data, *in vivo* HR-pQCT images in humans and micro-CT images in mice. These experiments indicate the portion of bone remodelling that can be attributed purely to mechanics and establish a baseline for futures studies evaluating mechanoregulation in patients.

Importantly, a combined *in vivo* and *in silico* validation, as shown in this study, has not yet been carried out. As such, algorithmic performance quantification was able to be carried out in in human radius geometries and mice. Previous studies provided validation using *in vivo* mouse loading experiments (Christen et al., 2012). However, this did not enable the demonstration of algorithmic functionality for load directions other than uniaxial compression, such as those observed in the human distal radius. The consistent results between our *in silico* and *in vivo* loading experiments indicate the validity of the MR algorithmic assumptions under diverse loading conditions. Corroborating the necessity for algorithmic validation in all six degrees of freedom, our *in silico* exerpiments identified possible performance deficits, when applied to complex loading regimes. Although MR’s performance was excellent for simulated adaptation, *in vivo* bone remodelling is not purely load-driven. Processes such as calcium homeostasis, wherein random bone remodelling may occur, will reduce MR’s performance. However, the findings of our mechanoregulation analysis reveal that the strain patterns overlap with the pattern observed by natural bone remodelling activity and can be used to estimate *in vivo* loading through our MR reverse engineering approach.

Predicted *in vivo* loading patterns in the mouse model were in good agreement with a previous study (Christen et al., 2012). Compared to the dataset used by Christen and colleagues (2012b), our LH results showed slightly larger moments while MR predictions were overall in good agreement with the previous study. Our LH results suggest a sizeable torsional component was induced in the caudal vertebra during daily activity, conflicting with the fact that the intervertebral discs limit the transmission of axial moments. This may be an artefact resulting from the homogeneous strain simplification of LH as the torsional moment was not detected using MR. LH predicted *in vivo* forces in the distal radius model were consistent with a previous study (Christen et al., 2013); however, predicted moments varied by order of magnitude. Christen et al., (2013) used layers of soft-tissue padding at the proximal and distal ends, which may have resulted in further homogenization of the strains throughout the radius. As such, this step may have limited the transmission of moment load at the interface between calcified- and soft-tissue. When comparing our results with a cadaveric study investigating distal radius load during various wrist motions (Smith et al., 2018), we find similar load-to-moment ratios indicating that additional padding may lead to an overestimation of momentum load. Our data also suggests that estimates in the distal radius may vastly vary from patient to patient. Despite the variance, an increase in loading was associated with increases in grip strength among patients. Such a relationship has been previously reported in cadaveric studies correlating grip strength with joint forces. In agreement with our results, models showed that approximately 26.3 N of force needs to be transmitted through the radius to obtain 10 N of grip strength (Putnam et al., 2000). Although this correlation was significant for loads estimated by MR and LH in the current study, this relationship was largely driven by single individuals with high grip strength. For future distal radius studies, grip strength should be considered as an inclusion parameter. Overall, our results indicate that the internal loads estimated by MR and LH are in good agreement with previous studies and can be linked to external factors such as grip strength in patients.

The principal algorithmic differences between approaches establish different future applications for LH and MR. MR prioritises remodelling sites, which are derived from two subsequent time-lapsed images. Accordingly, MR’s estimation is limited to the time frame between scans. LH estimates loading based on the bone morphology and is therby a cumulative estimate of all prior loading (load history). Loading during immobilisation treatment (Lill et al., 2003; Clayton et al., 2009; Spanswick et al., 2021), exercise (Troy et al., 2020), or loading interventions (Hughes et al., 2018), may differ from a patient’s load history, which is defined by every day and occupational activities. Thus, cumulative estimates of LH may be biased by the initial conditions. We showed that initial calibration of LH tends to improve differentiation between loading scenarios; however, this does not allow LH to achieve the same performance as MR. For the mouse loading experiment, this was evident in the delayed detection of significant differences between the loaded and control groups using LH compared to MR. For the present distal radius dataset, patients were skeletally mature adults and did not participate in a specific loading intervention. As a result, there was good agreement between MR and LH estimates. Note that the intact, contralateral radii used in the present study were taken from a patient cohort who had experienced a distal radius fracture. As such, loading the in the unfractured arm may have increased, particularly in cases involving fracture of the dominant arm. The resulting change in day-to-day loading may explain slightly higher predictions of MR compared to LH throughout the study. While our results indicate that MR is more sensitive to changes in loading, the algorithm is also more affected by imaging bias than LH. By utilising two subsequent HR-pQCT images, MR is subject to higher noise levels, movement artefacts and registration errors compared to LH (MacNeil and Boyd, 2008; Sode et al., 2011). Consequently, LH may be more suited to mouse studies, which can assess lifetime changes, but not for the timeframe of most clinical studies of antiresorptive therapies that often assess changes in BMD over a study duration of less than 2 years (Chen and Sambrook, 2012). Overall, our results have confirmed MR and LH’s capabilities for various applications using well-defined *in silico* loading and controlled experimental conditions. Accordingly, MR should be used when investigating designated time intervals in a longitudinal analysis and LH to assess the loading history in a cross-sectional fashion.

To quantify mechanoregulation, we have used a correct classification rate similar to the approach described by Tourolle né Betts et al. (2020). Here we show that by using the boundary condition derived by MR, we achieve significantly higher CCR values than LH for simulated, physiological, and extra physiological loading. Further, our results indicate that these differences are more pronounced when an extra physiological load was induced. Our results also show that using the simplified compressive boundary condition may be an acceptable choice when investigating trabecular bone mechanoregulation the healthy human distal radius. However,Johnson and Troy (2018) have shown that this simplified compression boundary condition may alter cortical and trabecular loading sharing. Therefore, the authors caution that such a simplified boundary condition may not be adequate for future studies investigating cortical and trabecular bone mechanoregulation. Although our results indicate a higher trend in CCR for loads estimated by MR, we cannot entirely rule out the possibility that inherent parallels between mechanoregulation analysis and MR synthetically inflate CCR within human distal radius data. However, our analysis of an *in vivo* loading model has provided experimental ground truth showing that estimations by MR reflected experimental conditions properly in mice. Further, our *in silico* validation showed that MR is highly sensitive, specific and accurate. Overall, our results indicate that mechanoregulation tends to be higher when analysing physiological loading derived by MR and thrives on a wealth of extra-physiological loading. Interestingly, our results also show that simple compression is an adequate simplification for the *in vivo* loading environment in the distal radius considering current limitations. Further, the results of our mechanoregulation analysis revealed a pronounced positive correlation between bone resorption and low strains for our mouse and a human model. This is in agreement with a previous study by Tourolle né Betts et al. (2020) investigating mechanoregulation in a rodent femoreal defect model, which indicated that mechanoregulated bone resorption mainly occurred within the distal and proximal fragments early during recovery. This relationship would indicate that osteoclastic activity may be more sensitive to local strain, and mechanoregulation may differ locally throughout the bone.

The proposed MR algorithm is subject to several limitations attributable to model assumptions as well as experimental and computational constraints. Regarding the animal experiments, the adjacent vertebra’s pinning procedure is limited in precision, and vibrations during the vertebra loading may create slight variations in loading direction and explain the observed higher variability in lateral bending. However, our results are comparable to a previous study (Christen et al., 2012) and represent the experimental setup sufficiently to provide validation for MR and LH. Regarding computational aspects, the method used to determine remodelling sites may include artefacts from scanning, such as beam hardening, motion artifacts, and partial volume effects or numerical inaccuracies of the image registration. However, *in vivo* micro-CT and HRp-QCT have been shown to have sufficient reproducibility for longitudinal bone structure assessment (Ellouz et al., 2014; Scheuren et al., 2020a). The FE model used was linear regarding material and geometry. Further, load cases were scaled and superimposed linearly during the optimisation to model the compounded loading effect. These simplifications would not capture any non-linear behaviour or visco-elastic effects; however, only small linear-elastic deformations are expected to occur during day-to-day activity. Future studies may expand this model with increasing computational power and investigate non-linear effects above yield strength that lead to bone failure (Schwiedrzik and Zysset, 2015).

## Conclusion

In conclusion, we have shown that MR is an enhanced load estimation algorithm tailored for longitudinal bone remodelling studies, achieving high sensitivity, specificity and accuracy by employing acknowledged mechanoregulation principles. The combined *in silico* and *in vivo* validation approach presented in this study proved to be a powerful benchmarking tool for the development of time-lapsed bone imaging analysis methods. Moreover, our results indicate that future studies may use grip strength as a functional surrogate for validation of estimated patient-specific physiological distal radius loads. Finally, our mechanoregulation analysis revealed considerable amounts of mechanically driven remodelling activity driven in human bone that may enable future studies to understand osteodegenrative disease.

## Author Contributions

**Authors’ Roles: Study design**: MW, NO, RM, CC. **Study conduct**: MW, FM. **Data collection**: MW. **Data analysis**: MW, NO, FM. **Data interpretation**: MW, FM, NO, RM, CC. **Drafting manuscript**: MW, CC. **Revising manuscript content**: MW, FM, NO, RM, CC. **Approving final version of manuscript**: MW, FM, NO, RM, CC. CC takes responsibility for the **integrity of the data analysis**.

## Acknowledgments

The authors acknowledge all participants who donated their time to participate in the study; the help of the study team at the University of Innsbruck for patient recruitment and data collection, specifically Dr. Lukas Horling, Dr. Kerstin Stock, Dr. Stefan Benedikt, and Katharina Gunther; the supervisory contributions of Dr. Patrik Christen and Prof. Dr. Michael Blauth in the clinical study; the contributions of Dr. Penny Atkins for efforts towards image registration; and computing ressources provided by the Swiss National Supercomputing Centre (CSCS).

## 1 Data Availability Statement

Data and Code will be made available upon reasonable request.

